# Fluctuating Selection among Individuals (FSI) as a Novel Genetic Drift in Molecular Evolution

**DOI:** 10.1101/2025.11.10.687654

**Authors:** Xun Gu

## Abstract

Fluctuating selection among individuals (FSI) refers to a mutation that exhibits different fitness effects among those individuals carrying this mutation, whereas the fitness of wildtype remains a constant. Consequently, the selection coefficient of a mutation should be interpreted by the means of the population average: 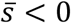 for deleterious, 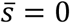 for neutral and 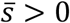 beneficial. For instance, a neutral mutation on average could be slightly deleterious in some individuals, as well as slightly beneficial in others. In this article, we developed a Wright-Fisher model under FSI and showed that the effect of FSI, measured by the FSI-coefficient 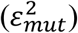, is important in molecular evolution especially when the effective population size (*N*_*e*_) is not very small. An intriguingly novel pattern of molecular evolution coined ‘selection duality’ emerges, that is, slightly beneficial mutations that satisfy 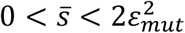 are subject to a negative selection such that the substitution rate *λ* less than the mutation rate *v*, revealing an *λ*∼ln*N*_*e*_ inverse relationship. Empirical analysis suggested, roughly, 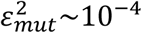, challenging the current wisdom of slightly deleterious mutations in molecular evolution. While FSI can be considered as new resource of genetic drift, an immediate question is to what extent this FSI-genetic drift would overperform the conventional *N*_*e*_-genetic drift. We thus developed a statistical method to predict the relative FSI strength 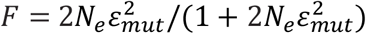 of each species. Our genome-wide study showed *F* > 0.5 in most metazoan species, suggesting that the FSI-genetic drift is, likely, dominant. Roughly, one may anticipate that *N*_*e*_-genetic drift could play a major role when *N*_*e*_<1000. The FSI theory may shed some lights on resolving the long-term neutralist-selectionist debate.

## Introduction

Galtier (2024) provided a historical perspective of the neutralist-selectionist debate in molecular evolution (Kimura 1968; Kern and Hahn 2018; Jensen et al. 2019), and highlighted how the debate has evolved from its origins to influence various aspects of modern evolutionary biology. Meanwhile, de Jong et al. (2024) evaluated the validity of both neutralist and selectionist arguments, finding that many of the tested predictions were sensitive to the underlying assumptions and can be consistent with both viewpoints. Thus, this central debate has casted some doubts whether the current wisdom of population genetics that underlies molecular evolution is theoretically sufficient; a recently-growing consensus between neutralists and selectionists. It has been indeed highly desirable to perform existed model examination and new model development (Hahn 2008; Hughes 2008; Wagner 2008; Nei et al. 2010; Munoz-Gomez et al. 2021; Gu 2021).

In population genetics, fixation models attempt to describe how a new mutation can spread in a population, ultimately toward being fixed rather than being lost, due to either selection or drift (Kimura 1962; 1983); one may see McCandlish and Stoltzfus (2014) for a comprehensive review. An implicit assumption of the fixation model is that any mutation has the same fitness among those individuals with the same genotype (homozygotes or heterozygotes). That said, a neutral mutation is selectively neutral for all individuals who carry the mutation, and so forth a deleterious or beneficial mutation. It should be noticed that human genetics revealed that mutations frequently exhibit different effects on individuals, challenging the fixed view of the mutational effects. In some studies, those effects were attributed to modifier genes (Nadeau 2001; Riodan and Nadeau 2017). For instance, incomplete penetrance occurs when a mutation shows deleterious effects in some individuals but not others; indeed, not all individuals with the same mutation show the disease phenotype (Riordan and Nadeau 2017; Eldar et al. 2009; Raj et al. 2010; Jensen et al. 2025). Moreover, Eldar et al. (2009) found the effect of partial penetrance on developmental evolution in bacteria.

In particular, the fluctuating selection among individuals (FSI) refers to any mutation that exhibits different fitness effects among individuals. In other words, the selection nature of a mutation should be interpreted by the means of the population average. For instance, one may anticipate that a neutral mutation could be slightly deleterious in some individuals, and slightly beneficial in others, but, on average, it is neutral. The population genetics of fluctuating selection between generations (FSG) has been extensively studied around 1970’s (Karlin and Lerikson 1974; Karlin and Lieberman 1974); one may see Gu (2024) for a recent commentary, but the issue of FSI was largely unexplored. The reason is mainly due to the presumption that the effect of FSI must be negligible in molecular evolution, as nearly-neutral mutations (slightly deleterious or beneficial) have played a major role on shaping the pattern.

Gu (2021) studied this FSI-problem, demonstrating that the effect of FSI is important in molecular evolution especially when the effective population size (*N*_*e*_) is not small. Yet, the model of FSI in Gu (2021) was found inaccurate. Recently, Gu (2025) addressed this issue, and conducted an extensive population genetics analysis to evaluate the effects of FSI on sequence divergence between species and genetic diversity within population. Intriguingly, a novel pattern of molecular evolution called ‘selection duality’ emerges: mutations that are statistically slightly beneficial are subject to negative selection, revealing a slow pace of molecular evolution in large population, and a signal of positive selection by the MK test simultaneously. Note that the former is the landmark evidence for the neutralist-view, whereas the latter is that for the selectionist-view. The condition for the emergence of selection-duality, intuitively speaking, is that the degree of selective advantage must be less than the effect of FSI. It means that in the case of no FSI, the selection-duality would be reduced to the strict neutrality.

Overall, it is theoretically plausible that FSI sheds some lights on the long-term neutralist-selectionist debate. An immediate question, apparently, is how to calculate the relative effect of FSI to the classical *N*_*e*_-genetic drift, which determines the range broadness of the selection duality. In the case of no FSI or *N*_*e*_ is very small, selection-duality would vanish as it is reduced to a single point of strict neutrality.

In this article, I formulate an FSI-model of molecular evolution that established an inverse relationship between the ratio of substitution rate to mutation rate (*λ*/*v*) and the logarithmic effective population size (ln *N*_*e*_). Compared with the classical nearly-neutral models of Ohta (1973), the FSI-model have three advantages: (*i*) it covers selection-duality mutations and deleterious mutations; (*ii*) it considers the effect of invariable sites; and (*iii*) it includes a non-zero rate component in a very large population. After using the conventional *d*_*N*_/*d*_*S*_ (the ratio of nonsynonymous rate to synonymous rate) as a proxy of *λ*/*v*, and *π*_*S*_ (the genetic diversity of synonymous mutations) as a measure of *N*_*e*_, we formulated the *d*_*N*_/*d*_*S*_∼ ln *π*_*S*_ analysis across a number of species, based upon which one can predict the strength of FSI for each species. Our results are exemplified by a genome-wide *d*_*N*_/*d*_*S*_∼ ln *π*_*S*_ analysis in metazoans and the relative importance of FSI-genetic drift to the *N*_*e*_-genetic drift in molecular evolution is discussed.

## Methods

### The two-allele haploid model of FSI

Consider a simple two-allele haploid system. Let *a* and *A* be the mutant and wild-type alleles at a particular locus, respectively, and *x* be the frequency of mutant *a*. Let *Z*_*A*_ or *Z*_*a*_ be the number of chromosomes produced by one parent *A*-chromosome or *a*-chromosome, respectively. It follows that the selection coefficient of allele *a* is simply defined by *s* = *Z*_*a*_/*Z*_*A*_-1. The FSI model postulates that *Z*_*a*_ is random variable among individual *a*-chromosomes with a variance denoted by *V*(*Z*_*a*_), whereas *Z*_*A*_ remains a constant. Thus, under the FSI model the variance of the selection coefficient is given by 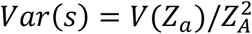.

Let *z*_*a*_ be the population mean of *Z*_*a*_ in a finite population. It appears that the population mean of selection coefficient of mutation-*a* is given by 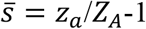. When the random fluctuation in selective coefficient is mainly due to the variation in genetic or epigenetic backgrounds that might be individual-specific, the effect of FSI on the change of allele frequency in each generation is determined by the variance of 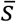, denoted by 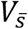 (Ohta 1973). Since the number of *a*-chromosomes is expected to be 2*Nx*, where *x* is the frequency of allele-*a* and *N* is the census population size, we have

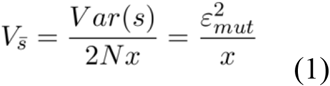

where 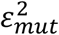 is called the FSI-coefficient; a large value means a strong FSI and *vice versa*; the subscript indicates that this FSI is induced by the mutation. It should be noticed that Ohta (1972) speculated that 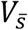 should inversely related to the allele frequency under FSI, making a striking distinction from the fluctuating selection between generations as driven by seasonal environmental changes.

### The Wright-Fisher model under FSI

Consider a random mating population of a monoecioys diploid organism. In a finite population, an individual produces a huge number of offspring and that exactly *N*_*e*_ of those survive to maturity. Let *a* and *A* be the mutant and wild-type alleles at a particular locus, respectively, whose fitness effects are additive. The FSI model postulates that the fitness effect of mutant *a* is fluctuating among individuals, whereas that of wild-type *A* remains a constant. Therefore, the relative fitness of genotype *AA, Aa*, or *aa* is, on average, given by 1, 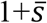 and 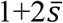, respectively; 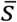 is the mean selection coefficient (*s*) of mutant *a*.

The amount of change (*Δx*) of gene frequency per generation (where *Δt* is the generation time) under FSI can be decomposed into two independent components, that is

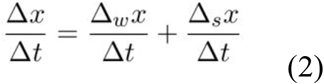

where *Δ*_*w*_*x* and *Δ*_*s*_*x* are the frequency changes caused by the Wright’s finite sampling and FSI, respectively. Under the Wright-Fisher model, the first and second moments of *Δ*_*w*_*x* are given by

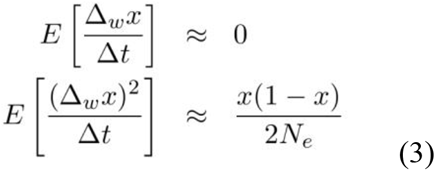

respectively, where *N*_*e*_ is the effective population size. Next the expected gene frequency change (per generation) with respect to FSI can be derived as follows

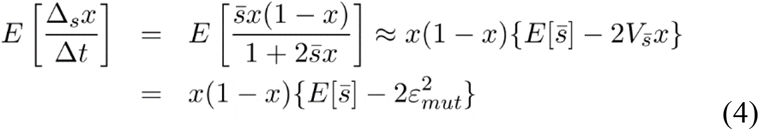

The Taylor’s expansion in Eq.(4) was carried out with respect to 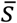, up to 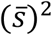, where the absolute value of 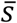 is much less than one. The last equation of Eq.(4) used the relationship 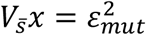 implied by Eq.(1). In the same manner, the second moment of *Δ*_*s*_*x* is given by

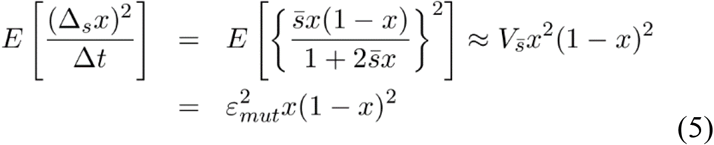

#### The diffusion-limit approximation of FSI model

It is mathematically convenient to apply the diffusion approximation to study the Wright-Fisher model under FSI. Intuitively, the infinitesimal mean *μ*(*x*) can be approximated by 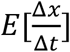, and the infinitesimal variance 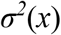 by 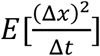. Together with Eqs.(2)-(5), we obtain

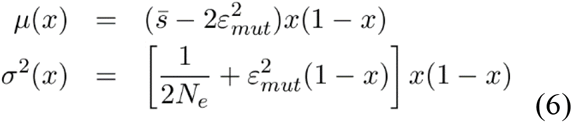

To be concise, we use 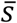 to replace 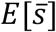 in Eq.(6). While *μ*(*x*) describes how determinative factors shape the mean frequency change, *σ*^*2*^(*x*) describes the random effects of genetic drifts. Noticeably, FSI may have profound impacts on both selection and genetic drift. The FSI-coefficient 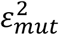 not only negatively affects the mean change of gene frequency, but also provides a novel resource of genetic drift in molecular evolution, called the FSI-drift. One may further anticipate that the FSI-drift would be dominant in a large population, whereas the *N*_*e*_-drift becomes dominant in a small population.

## Results

### Fixation probability and the rate of nucleotide substitution under FSI

The rate of nucleotide substitution (*λ*) plays a central role in the study of molecular evolution. Under fixation models, mutations are usually described as random events that have a rate of occurrence (mutation rate) and a contribution to fitness (selection). Suppose we have a diploid population of size *N* (the census population size) and we assume that mutations appear with a rate of *v*. Then the total rate that such mutations appear in the population per generation is given by 2*Nv*. Since only a fraction of these mutations will eventually go to fixation, to calculate the rate of molecular evolution (λ), we need to multiply 2*Nv* with the fixation probability of a new mutation (Kimura 1962). Let *u*(*p*) be the fixation probability of a mutation in the population, with the initial frequency *p*. Then, the formula of substitution rate can be concisely written as follows

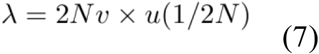

where the initial frequency set by 1/2*N* represents rare, single *de novo* mutation event.

It appears that derivation of the fixation probability *u*(*p*) is the key step to solve the problem of substitution rate. To this end we define two variables at first. One is the selection-FSI ratio *ρ*, the (two-fold) ratio of 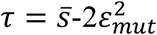 to the FSI-coefficient 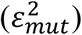, that is,

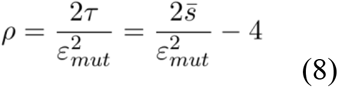

Intuitively speaking, the selection-FSI ratio *ρ* measures the relative magnitude of the selection pressure to the FSI strength.

Meanwhile, FSI emerges as a new resource of genetic drift, which can be measured by the FSI-strength, 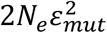. Reading the FSI-strength by 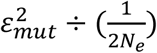, we thus claim a dominant FSI-drift when 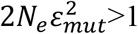, or a dominant *N*_*e*_-drift when 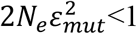. In this sense, one may define a relative measure of FSI-strength (*F*) by

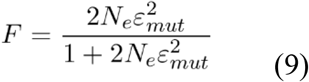

Apparently, *F* increases from 0 at 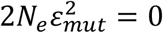 (no-FSI), which approaches to 1 when 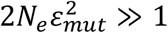.

The standard procedure for deriving *u*(*p*) under the diffusion model starts from the calculation of *G*(*x*), that is,

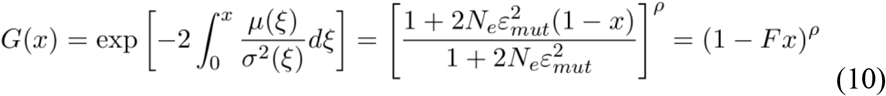

It follows that the fixation probability for a given initial frequency *p* can be calculated by

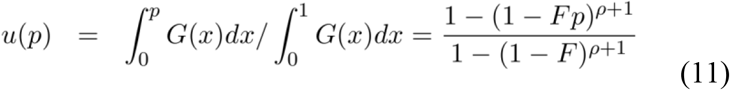

Under the approximation of very small initial frequency (*p* → 0), the fixation probability of Eq.(11) can be further simplified by

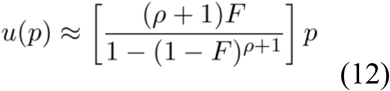

Together with Eq.(7) and (12), we obtain the formular of substitute rate (*λ*) as follows

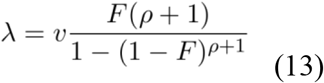

It appears that, in addition to the mutations rate (*v*), the substitution rate depends on two parameters: the selection-FSI ratio (*ρ*), and the relative FSI-strength (*F*). As illustrated by Fig.1, the effects of FSI on the substitution rate can be concisely summarized as follows.

**Fig.1.**
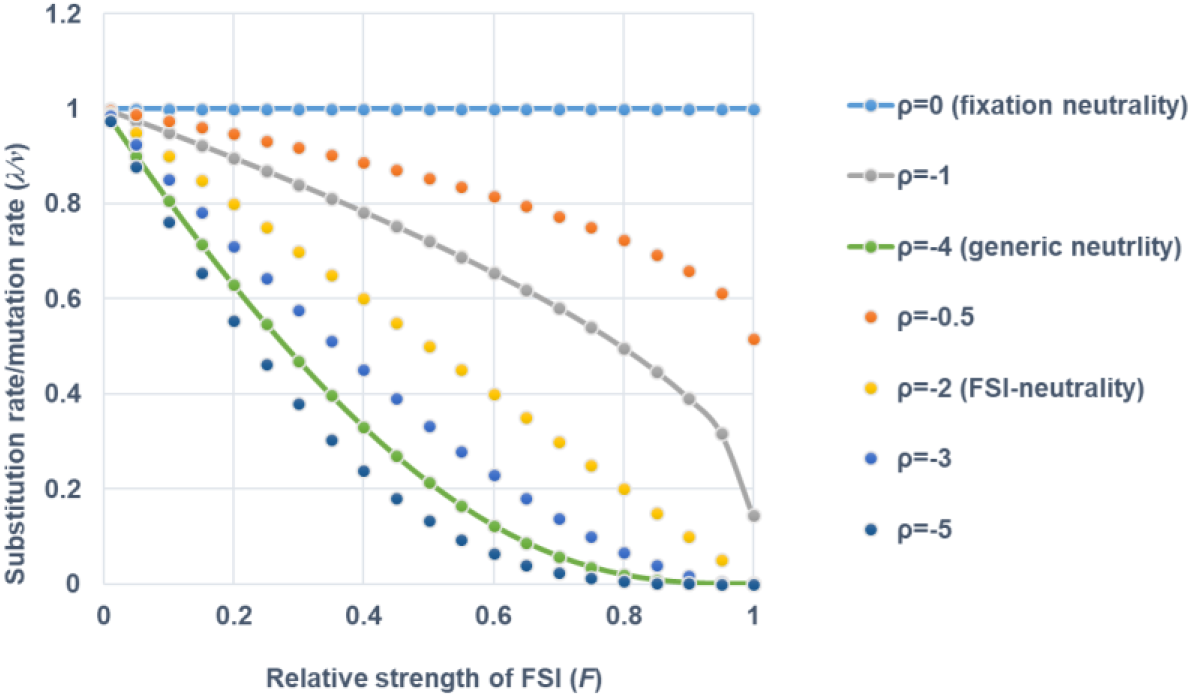
The impact of FSI on the substitution rate in the case of no adaptive evolution (*ρ* ≤ 0). Selection-duality is defined by the region between the solid line of fixation neutrality (*ρ*=0) and that of generic neutrality (*ρ*=-4). The Ω-region is defined by that between the solid line of fixation neutrality (*ρ*=0) and that of *ρ*=-1), in which the substitution rate is nonzero even in a very large population.

i. The substitution rate equals to the mutations rate (*λ* = *v*) when *ρ* = 0; as well as *λ* > *v* when *ρ* > 0, or *λ* < *v* when *ρ* < 0, respectively.
ii. When *ρ* > 0, the substitution rate (*λ*) increases with the increasing of *F* or the effective population size (*N*_*e*_).
iii. When *ρ* < 0, the substitution rate (*λ*) decreases with the increasing of *F* or the effective population size (*N*_*e*_).
iv. As *N*_*e*_ → ∞, the substitution rate (*λ*) approaches to

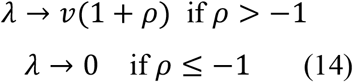

In other words, in a very large population, the substitution rate approaches to a nonzero constant that is less than the mutation rate when 0 > *ρ* > −1, a new mechanism for molecular evolution in a large population (Gillespie 2000, 2001; Lefear et al. 2014).
v. In the case of no FSI, i.e., *ε*_*mut*_^*2*^=0, one can verify that 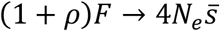, and 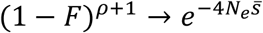, resulting in the classical formula (Kimura 1962), i.e.,

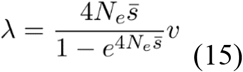

In short, the substitution rate-mutation rate ratio (*λ*/*v*) normally determined by *ρ* and *F* under FSI, but both of which would merge into a single variable 4*N*_*e*_*s* in the case of no-FSI.

### Emergence of selection-duality

Noticeably, Eq.(13) shows that when 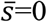 under FSI so that *ρ* = −4, the substitution rate (*λ*) is always less than the mutation rate (*v*). That is, a generically neutral mutation 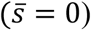 could be subjecting to a negative selection. On the other hand, a mutation reveals a neutral evolution (*λ* = *v*) when *ρ* = 0, or 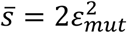, highlighting that this mutation should be, on average, slightly beneficial. Those two facts are challenging the traditional wisdom of neutrality. To further clarify this issue, one may call the case of *ρ* = 0 (or 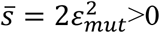) *the substitution neutrality*, corresponding to the neutrality in molecular evolution: the substitution rate equals to the mutation rate (*λ* = *v*). Meanwhile, *the generic neutrality* refers to the case of *ρ* = −4 (or 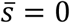), corresponding to the neutrality on average among individuals in the population.

An intriguing phenomenon called *selection-duality* then emerges between the generic neutrality and the substitution neutrality: a slightly beneficial mutation defined by 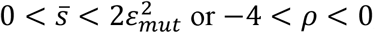 or −4 < *ρ* < 0, is subjecting to a negative selection such that the substitution rate is less than the mutation rate (*λ* < *v*). The arising of selection-duality is mainly due to the statistical nature of FSI. Consider a normal-like distribution of FSI that can be divided into the negative side (*s*<0) and the positive side (*s*>0), respectively. It has been known that the effect of negative side of selection coefficient to reduce the fixation probability is more severe than that of positive side to increase the fixation probability. Consequently, generic neutrality at 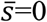 implies an equal proportion of both sides, revealing an overall negative selection. Under the substitution neutrality, the proportion of positive side is slightly greater than that of negative side such that the net effect on the fixation probability has been perfectly compromised.

While the generic neutrality and the substitution neutrality define the boundary of selection-duality, Gu (2025) pointed out that the middle-point of selection-duality (at 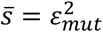 or *ρ* = −2), coined the FSI-neutrality, may play a pivotal role in population genetics, as it can serve as an appropriate null hypothesis for population genetics tests under FSI.

### The rate-In[*N*_*e*_] inverse relationship and the *d*_*N*_/*d*_*S*_∼ In*π*_*S*_ analysis

The inverse relationship between the evolutionary rate (λ) and the effective population size (*N*_*e*_) has become the landmark prediction of the nearly-neutral theory (Ohta 1973). Two decades later, Ohta (1993) tested this inverse relationship by using the rate ratio of nonsynonymous to synonymous substitutions (*d*_*N*_/*d*_*S*_) of encoding genes, a proxy of the ratio of substitution rate to the mutation rate (*λ*/*v*). Based on a limited number of genes, the author showed a roughly two-fold of the mean *d*_*N*_/*d*_*S*_ in the human lineage as that in the mouse lineage. Since the effective population size of humans is much smaller than that of mice, it has been widely cited as the textbook evidence to support the nearly-neutral theory (Li 1997; Lynch 2006; Lynch et al. 2011; Kosiol et al. 2008; Ellegren 2009). However, this *λ* − *N*_*e*_ inverse relationship has suffered a severe scale-problem (Ellegren 2009; Galtier et al. 2016; Gu 2021). For instance, Galtier et al. (2016) reported a genome-wide analysis in metazoans. Using the synonymous genetic diversity (*π*_*S*_) as a proxy of *N*_*e*_, they showed that the genome-wide *d*_*N*_/*d*_*S*_ ratio is negatively correlated with the log of *π*_*S*_. In other words, molecular evolution indeed evolves slowly in a large population, but still considerably faster than the prediction from the nearly-neutral theory.

Eq.(13) implies that FSI may provide a solution to this scale-discrepancy. To this end, one should take the heterogeneity of selection pressures at different nonsynonymous nucleotide sites into account. Suppose that the substitution rate at a nonsynonymous nucleotide site follows Eq.(13) under FSI. Without adaptive evolution the selection-FSI ratio is always negative, i.e., −∞ < *ρ* < 0. We further assume that the FSI-coefficient 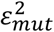 is the same across all nonsynonymous sites. Hence, the *ρ-*variation (−∞ < *ρ* < 0) reflects the variation of the mean selection coefficient 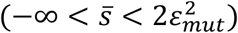 among nonsynonymous sites. Note that the substitution rate of gene (*λ*_*g*_) is always less than the mutation rate because of *ρ* < 0.

We implement the Ohta-Gu model for the *ρ-*variation among nonsynonymous sites. A proportion (*θ*_*inv*_) of nonsynonymous sites are invariable, at which all mutations are lethal. For the rest nonsynonymous sites that are variable, rewrite *ρ* = −*z*|*ρ*_*g*_|, where |*ρ*_*g*_| is the absolute mean *ρ* over variable nonsynonymous sites, and *z* > 0. It is further assumed that z is a random variable that follows an exponential distribution. It appears that the Ohta model (Ohta 1973) is a special case when *θ*_*inv*_=0. From Eq.(13), the mean substitution rate of a gene is then given by

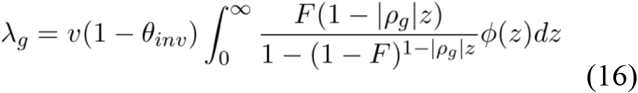

As shown by the Appendix in details, the substitution rate of Eq.(16) can be approximated by

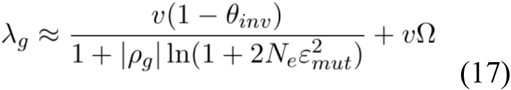

Intuitively speaking, Eq.(17) claims that the substitution rate can be divided two components. The first one is *N*_*e*_-dependent: the log-inverse relationship *λ*_*g*_∼ ln *N*_*e*_ holds when 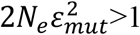, or the inverse relationship *λ*_*g*_∼*N*_*e*_ holds when 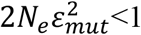. The second term represents the substitution rate in a very large population, where Ω is the expected proportion of nucleotide sites in the range of −1 ≤ ρ < 0. Note that Eq.(17) implies that the notion of molecular clock (Zuckerkandl and Pauling 1965) may vary locally due to the variation of effective population size but the scale of variation tends to be logarithmic.

We apply Eq.(17) to analyze nonsynonymous substitutions in encoding genes. Let *d*_*N*_/*d*_*S*_ be the ratio of nonsynonymous distance (*d*_*N*_) to synonymous distance (*d*_*S*_) of an encoding gene between two closely-related species, and *π*_*S*_ be the genetic diversity of synonymous mutations in the population. Following the common practice, the *d*_*N*_/*d*_*S*_ ratio can be used as a proxy of the ratio of the (nonsynonymous) substitution rate to the mutation rate (*λ*_*g*_/*v*). Meanwhile, the synonymous diversity *π*_*S*_ is used as a measure of effective population size (*N*_*e*_) by the formula *π*_*S*_ = 4*N*_*e*_*v*. Both statements are based upon the assumption that synonymous mutations are strictly neutral. In the case when FSI at nonsynonymous sites is strong, the parameter Ω is close enough to its limit 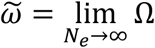 that is *N*_*e*_-independent. Replacing *λ*_*g*_/*v* by *d*_*N*_/*d*_*S*_, and *N*_*e*_ by *π*_*S*_/(4*v*), and Ω by 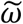 in Eq.(17), we obtain

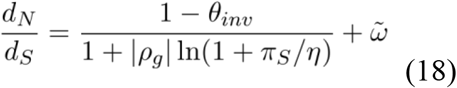

where 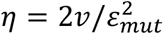, the (two-fold) ratio of mutation rate to the FSI coefficient at nonsynonymous sites. In other words, 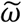 is the predicted *d*_*N*_/*d*_*S*_ ratio when the effective population size is very large.

### Computational procedure and prediction of FSI-strength

Suppose we have a certain number of species under study, called focus species. For each focus species, the *d*_*N*_/*d*_*S*_ ratio could be estimated based on a closely-related species, and the synonymous genetic diversity (*π*_*S*_) is obtained by population genetics screening. The main goal of *d*_*N*_/*d*_*S*_∼ ln *π*_*S*_ analysis is to statistically predict the relative FSI strength (*F*) or the (absolute) FSI strength 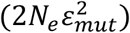 for each focus species. There are four unknown parameters in the non-linear *d*_*N*_/*d*_*S*_∼ ln *π*_*S*_ regression analysis across multiple species based on Eq.(18), i.e., *θ*_*inv*_, *η*, 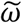, *ρ*_*g*_. Simulation analysis has shown that estimation of those parameters may be subject to a high level of statistical uncertainty except when the number of species is at least several hundreds. Nevertheless, under the assumption of strong FSI strength 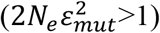, one can show that Eq.(18) can be well approximated by

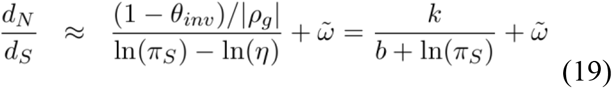

where *k* = (1 − *θ*_*inv*_)/|*ρ*_*g*_| and *b* = − ln *η*. It has been shown that the sampling variances of unknown parameters in Eq.(19) can be considerably reduced, compared to Eq.(18). A procedure in the *R*-package was implemented.

The final step of our *d*_*N*_/*d*_*S*_∼ ln *π*_*S*_ analysis is to predict the FSI strength of nonsynonymous sites for each species. Recall the definition of 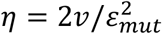. Thus, given the synonymous diversity (*π*_*S*_) for a given species, one can predict 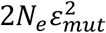 by

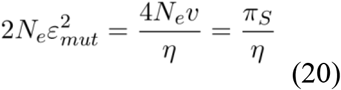

as long as *η* is known. Let 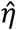 be the estimate of *η*, the relative strength of FSI for nonsynonymous sites is given by

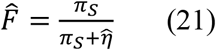

### Case study: molecular evolution of metazoans

Galtier et al. (2016) reported a genome-wide analysis in metazoans, a genome-wide set of protein-coding genes in 44 pairs including 18 vertebrate and 26 invertebrate pairs. Each pair consisted of two closely-related species to estimate the synonymous distances (*d*_*S*_) and the nonsynonymous distance (*d*_*N*_), respectively. Moreover, the synonymous genetic diversity (*π*_*S*_) in the focal species of each specie pair was used as a proxy of the effective population size. Fig.2 represents the plotting of *d*_*N*_/*d*_*S*_ to ln *π*_*S*_, showing a strong negative correlation between them; the straight line (*p*-value <0.001). This inverse *d*_*N*_/*d*_*S*_∼ ln *π*_*S*_ provides evidence for supporting the notion of FSI-driven molecular evolution when 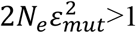. By fitting the data with Eq.(19) (Fig.2), we estimated the mutation-to-FSI ratio 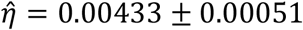.

**Fig.2.**
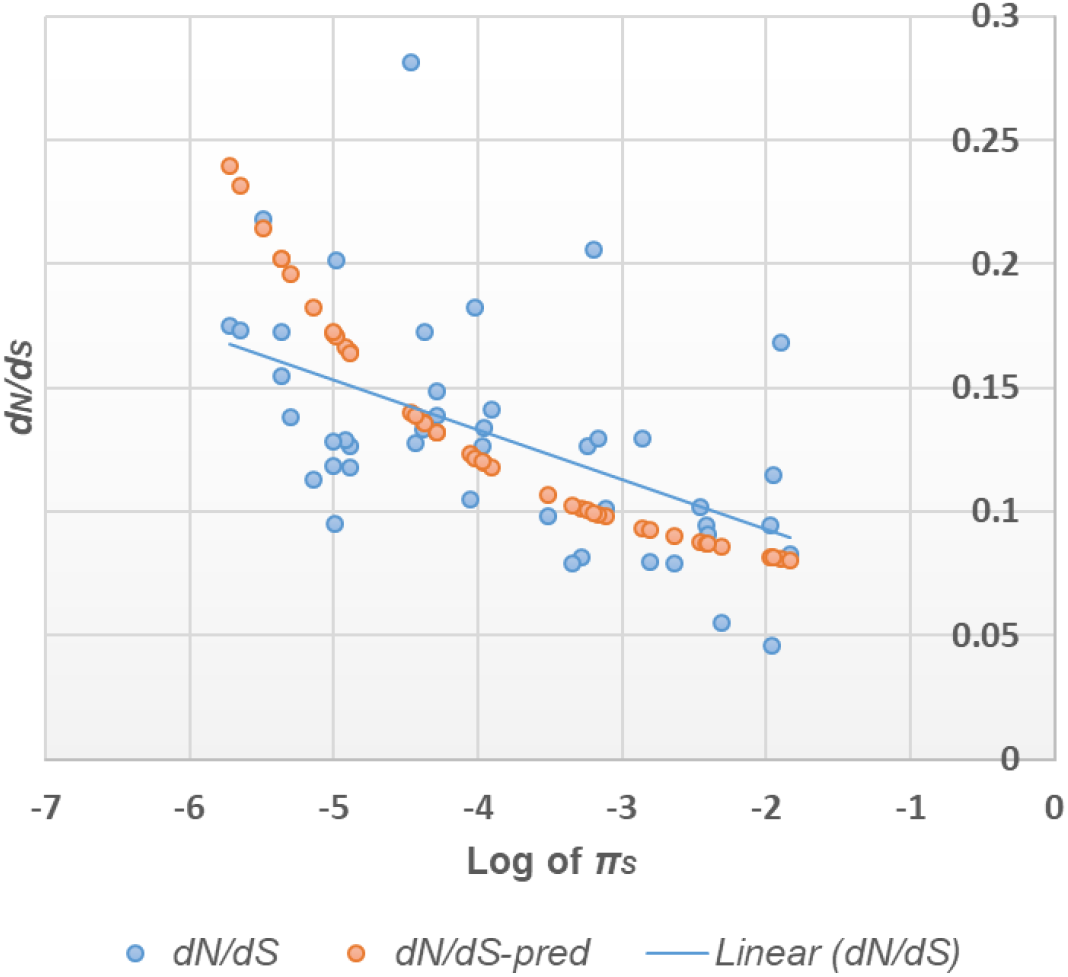
The potting of *d*_*N*_/*d*_*S*_ against the log of synonymous genetic diversity (*π*_*S*_). Data were from Galtier et al. (2016) including 18 vertebrate and 26 invertebrate pairs. See Table 1 for the name list of all species. Blue circles represent the observed *d*_*N*_/*d*_*S*_ and *π*_*S*_ of each species, whereas the orange circles represent the predicted *d*_*N*_/*d*_*S*_, based on the estimated model parameters and the observed *π*_*S*_.

Table 1 represents the list of predicted 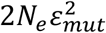 as well as *F* in 44 metazoans. Surprisingly, we found that 41 species show 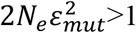 or *F*>0.5. Those species with *F* ≤ 0.5 are *Camponotus ligniperdus* (Carpet ant), *Reticulitermes flavipes* (Eastern subterranean termite), and *Homo sapiens* (Human). Our finding suggests that, during the long-term metazoan evolution, the FSI-coefficient 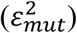 roughly remains a constant, whereas the effective population size varies in several magnitudes among species, resulting in up to three-fold changes of the substitution rate of genome evolution. We therefore conclude that FSI-genetic drift may play a more importance role than *N*_*e*_-genetic drift in molecular evolution of most metazoans.

**Table 1.** List of predicted FSI strength 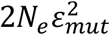 and the relative FSI strength (*F*) for each species.

| *Focal_species* | *Outgroup* | *Taxonomy* | $2N_e\epsilon_{mut}^2$ | $F$ |
| --- | --- | --- | --- | --- |
| *Allolobophora_chlorotica_L2* | *Aporrectodea_icterica* | Annelida | 36.29 | 0.973 |
| *Armadillidium_vulgare* | *Armadillidium_nasatum* | Arthropoda | 13.96 | 0.933 |
| *Artemia_franciscana* | *Artemia_sinica* | Arthropoda | 9.18 | 0.902 |
| *Camponotus_ligniperdus* | *Camponotus_aethiops* | Arthropoda | 0.80 | 0.444 |
| *Culex_pipiens* | *Culex_torrentium* | Arthropoda | 34.14 | 0.972 |
| *Halictus_scabiosae* | *Halictus_simplex* | Arthropoda | 1.67 | 0.626 |
| *Melitaea_cinxia* | *Melitaea_didyma* | Arthropoda | 34.67 | 0.972 |
| *Messor_barbarus* | *Messor_structor* | Arthropoda | 4.64 | 0.823 |
| *Necora_puber* | *Carcinus_aestuarii* | Arthropoda | 4.22 | 0.808 |
| *Reticulitermes_grassei* | *Reticulitermes_flavipes* | Arthropoda | 1.00 | 0.500 |
| *Thymelicus_lineola* | *Thymelicus_sylvestris* | Arthropoda | 17.42 | 0.946 |
| *Thymelicus_sylvestris* | *Thymelicus_lineola* | Arthropoda | 14.70 | 0.936 |
| *Eunicella_cavolinii* | *Eunicella_verrucosa* | Cnidaria | 9.49 | 0.905 |
| *Abatus_cordatus* | *Abatus_agassizi* | Echinodermata | 2.81 | 0.738 |
| *Echinocardium_cordatum_B2* | *Echinocardium_mediterraneum* | Echinodermata | 33.84 | 0.971 |
| *Echinocardium_mediterraneum* | *Echinocardium_cordatum_B2* | Echinodermata | 10.88 | 0.916 |
| *Ophioderma_longicauda_L1* | *Ophioderma_longicauda_L3* | Echinodermata | 9.96 | 0.909 |
| *Chlorocebus_aethiops* | *Macaca_mulatta* | Mammalia | 1.83 | 0.647 |
| *Eulemur_coronatus* | *Eulemur_mongoz* | Mammalia | 1.63 | 0.620 |
| *Eulemur_mongoz* | *Eulemur_coronatus* | Mammalia | 1.43 | 0.588 |
| *Galago_senegalensis* | *Nycticebus_coucang* | Mammalia | 1.78 | 0.640 |
| *Homo_sapiens* | *Pan_troglodytes* | Mammalia | 0.85 | 0.461 |
| *Lepus_granatensis* | *Lepus_americanus* | Mammalia | 3.35 | 0.770 |
| *Macaca_mulatta* | *Chlorocebus_aethiops* | Mammalia | 1.83 | 0.647 |
| *Microtus_arvalis* | *Microtus_glareolus* | Mammalia | 7.25 | 0.879 |
| *Nycticebus_coucang* | *Galago_senegalensis* | Mammalia | 1.21 | 0.548 |
| *Pan_troglodytes* | *Homo_sapiens* | Mammalia | 3.07 | 0.754 |
| *Propithecus_coquereli* | *Varecia_variegata_variegata* | Mammalia | 4.59 | 0.821 |
| *Varecia_variegata_variegata* | *Propithecus_coquereli* | Mammalia | 2.89 | 0.743 |
| *Crepidula_fornicata* | *Crepidula_plana* | Mollusca | 20.93 | 0.954 |
| *Mytilus_galloprovincialis* | *Mytilus_californianus* | Mollusca | 21.76 | 0.956 |
| *Ostrea_edulis* | *Ostrea_chilensis* | Mollusca | 4.38 | 0.814 |
| *Physa_acuta* | *Physa_gyrina* | Mollusca | 21.89 | 0.956 |
| *Sepia_officinalis* | *Sepiella_japonica* | Mollusca | 1.63 | 0.620 |
| *Caenorhabditis_brenneri* | *Caenorhabditis_sp.10* | Nematoda | 24.26 | 0.960 |
| *Lineus_longissimus* | *Lineus_ruber* | Nemertea | 10.26 | 0.911 |
| *Aptenodytes_patagonicus* | *Aptenodytes_forsteri* | Sauropsida | 3.04 | 0.753 |
| *Chelonoidis_nigra* | *Chelonoidis_carbonaria* | Sauropsida | 1.14 | 0.532 |
| *Emys_orbicularis* | *Trachemys_scripta* | Sauropsida | 3.34 | 0.770 |
| *Eudyptes_moseleyi* | *Pygoscelis_papua* | Sauropsida | 1.65 | 0.623 |
| *Parus_caeruleus* | *Parus_major* | Sauropsida | 4.92 | 0.831 |
| *Hippocampus_guttulatus* | *Hippocampus_kuda* | Teleostei | 1.14 | 0.532 |
| *Ciona_intestinalis_A* | *Ciona_intestinalis_B* | Urochordata | 8.62 | 0.896 |
| *Ciona_intestinalis_B* | *Ciona_intestinalis_A* | Urochordata | 38.90 | 0.975 |

An intriguing question is what is a typical value of 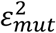. We estimated it roughly by two approaches. First, given the mutation rate *v* ≈ 3 × 10^−7^per generation, we obtain 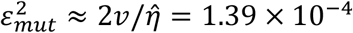. Second, the FSI strength 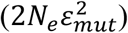 in humans is predicted to be 0.461 (Table 1). Given *N*_*e*_ = 5000 in the human population, we obtained 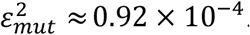. Tentatively, we believe that 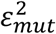 should be roughly at the magnitude of 10^−4^.

## Discussion

### The significant role of FSI-genetic drift in molecular evolution

The FSI (fluctuating selection among individuals) theory of molecular evolution is based upon the premise that fitness may fluctuate among individuals that carry the mutation. The relative importance of FSI-genetic drift to the conventional *N*_*e*_-genetic drift (due to finite population size) can be measured by the relative FSI strength (*F*), defined by Eq.(9): if *F* is close to 0, the *N*_*e*_-genetic drift is dominant; whereas the FSI-genetic drift is dominant if *F* is close to 1. Tentatively, one may set *F*=0.5 as an empirical criterion to weigh between those two genetic drifts. Table 1 showed that the relative FSI-strength *F* is over 0.5 in most metazoan species, suggesting that the FSI-genetic drift, rather than the *N*_*e*_-genetic drift, plays a major role in metazoan genome evolution.

Meanwhile, it should be noticed that the classical neutral or nearly-neutral theory applies when the effective population size is small. An immediate question is what is the cutoff in practice. Suppose that the FSI-coefficient is round 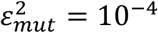. It appears that *F*=0.5 when the effective population size *N*_*e*_ = 5,000. So, roughly speaking, the FSI-genetic drift and the *N*_*e*_-genetic drift both play a role when *N*_*e*_ is around 5,000. The *N*_*e*_-genetic drift becomes dominant when *N*_*e*_ is less than 1,000, whereas the FSI-genetic drift becomes dominant when *N*_*e*_ is greater than 10,000. Interestingly, as the effective population size of humans is about 5,000, molecular evolution of nonsynonymous sites in human genes could be driven by both genetic drifts.

### Roles of selection-duality versus slightly deleterious mutations in molecular evolution

Somewhat unexpectedly, the theory of FSI toppled one of golden-standards of neutral theory: the substitution rate equals to the mutation rate (*λ* = *v*) if and only if the mutation is strictly neutral without any FSI. Consequently, the pattern selection-duality at nonsynonymous sites emerges: under FSI, slightly beneficial mutations may be subject to a negative selection, resulting in a substitution rate less than the mutations rate, as well as an inverse relationship between the substitution rate and the log-of effective population size.

To avoid confusion, here we define nearly neutral mutations by *λ* < *v*. Suppose the FSI coefficient to be 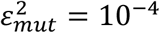. The FSI theory implies that there are two components in nearly neutral mutations. The first component is the selection-duality with the mean selection coefficient falling in the range of 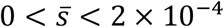 (slightly beneficial), and the second component is the slightly deleterious mutations 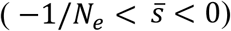. When the effective population size is small, say, *N*_*e*_ < 1000, selection duality mutations turn out to be virtually neutral, and slightly deleterious mutations may play a role in molecular evolution. On the other hand, when the effective population size is intermediate or large, say, *N*_*e*_ > 10000, the *N*_*e*_-drift barrier would hinder the fixation of those slightly deleterious mutations, and those selection-duality mutations would be driven by the dominant FSI-genetic drift. Overall, the FSI-theory may provide new insights on resolving the long-term neutralist-selectionist debate.

### Effects of background FSI

For the sake of simplicity, the FSI model we have developed assumes no-FSI in wildtype allele. One may wonder to what extent the main results of FSI theory could be affected by the existence of background FSI, or *FSI*_*bg*_, the stochastic variation of offspring number commonly for all individuals. While the detailed analysis will be published elsewhere, a concise summary is shown below. Let *FSI*_*mut*_ be the overall FSI-coefficient induced by the mutant. Under a more general framework, one can show that the FSI coefficient is simply given by 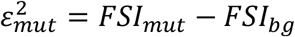. Hence, the pattern of FSI emerges if 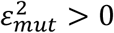, i.e., the level of mutant FSI is greater than that of the background FSI. In other words, the null hypothesis of no-FSI simply means that the mutant FSI equals to the background FSI, which would be integrated into the renormalized effective population size. It is therefore technically convenient to postulate that the mutant is subjecting to FSI, whereas the wildtype has no FSI. A final but intriguing question is what if *FSI*_*mut*_ < *FSI*_*bg*_, an emergence of anti-FSI such that a mutant is trying to reduce the background FSI of wildtype individuals. We shall further address this issue in the future study.

### Biological mechanisms of FSI: intrinsic fluctuation versus local adaptation

From the evolutionary view, the biological mechanisms of FSI can be roughly classified into two categories: the intrinsic randomness versus the local adaptation. In essence, the intrinsic randomness reflects the stochastic nature of genotype-phenotype mapping (Dowell et al. 2010; Lehner 2013; Binder et al. 2015), e.g., regulatory or epigenetic effects triggered by stochastic fluctuations in development stages (Ozbudak et al. 2002; Elowitz et al. 2002; Suel et 2007; Chang et al. 2008). Intrinsic randomness can arise through a variety of mechanisms, including epistasis (genetic interactions) (Mullis et al, 2018; Taylor and Ehrenreich 2014), genetic background (Chandler et al. 2013), or epigenetics (Maamar et al. 2007; Raj et al. 2008). In development, variable expressivity occurs when a mutation shows quantitatively different phenotypic effects among individuals (Vu et al. 2015). Differences in gene expression between affected and unaffected individuals were frequently used to explain the randomness of penetrance (Raj et al. 2010; Hume 2000; Wernet et al 2006).

Secondly, one may anticipate that FSI is intrinsically related to the evolution and maintenance of phenotypic plasticity that may imply local adaptation (Vu et al. 2015; Khourt et al. 1988; Sommer 2020; Lee et al. 2022; Wang et al. 2023). It usually involves the sophisticated genetic-environmental interactions underlying the so-called phenotype plasticity (Lai et al. 2024; Wang et al. 2023; Lee et al. 2022; Sommer et al. 2020), which possibly is shaped by the local adaptation (He et al. 2023). Though the detail may have been beyond the scope of current study, we pinpoint out that FSI could be the result of local adaptation for some individuals, but not necessary.

### Some technical issues

#### Ohta-Gu model versus Ohta-Kimura model

Our current study invokes the Ohta-Gu model: a proportion (*θ*_*inv*_) of nonsynonymous nucleotide sites are invariable, at which any mutation is assumed lethal, and at the rest of variable sites, the (negative) selection-FSI ratio (−*ρ*) of mutations follows an exponential distribution. Kimura (1979) argued that the exponential model (Ohta 1973) (with no invariable sites) was insufficient to account for the selection heterogeneity among mutations. Instead, he suggested a gamma distribution with the shape parameter γ=1/2; note that the exponential model is a special case with γ=1. It should be noticed that a less-than-one shaper parameter of the gamma distribution controls both portions of highly deleterious mutation and nearly neutral mutations, which is biologically unrealistic. Instead, the Ohta-Gu model allows a proportion of invariable sites to describe mutations that are highly deleterious or even lethal) is biologically more plausible.

#### Analogy to Kimura’s basic formula of molecular evolution

While we have shown that the classical formula of molecular evolution (Kimura 1962) is a special case of Eq.(13), it is desirable to have a form of formula that is intuitively analogous to the classical formula. To this end, we introduce a new variable *Q*, the FSI-selection intensity, by 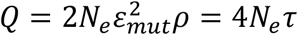, as well as the *f*-index of FSI by

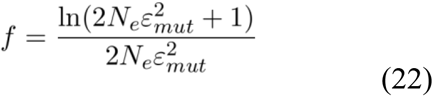

Putting together, the substitution rate of Eq.(13) can be written as follows

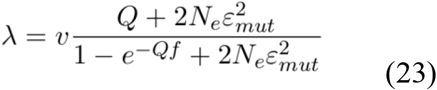

Intuitively speaking, in addition to the mutation rate (*v*), the substitution rate (*λ*) is determined by the FSI selection intensity (*Q*), the FSI-intensity 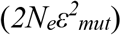, and the *f*-index of FSI. One can easily verify that, when *ε*^*2*^_*mut*_=0, we have *f*=1 and 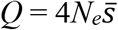 such that Eq.(23) would be reduced to the formula of Kimura (1962).

## Appendix: Derivation of the substitution rate of a gene

To simplify the notation, we rewrite *ρ* = −*z*|*ρ*_*g*_|, where |*ρ*_*g*_| is the absolute value of the mean of *ρ* over (nonsynonymous) sites, and *z* is a positive, random variable that follows an exponential distribution, with a mean of 1, that is,

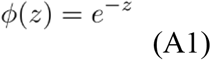

Under the Ohta-Gu model, the mean substitution rate of a gene is then calculated by

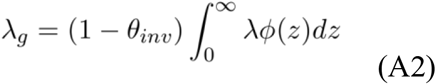

To obtain a close-form of *λ*_*g*_, we first notice that the substitution rate given by Eq.(6) has two distinct patterns when *ρ* < 0. As *F* increases to 0 to 1, λ decreases from *v* toward 0 in the case of *ρ* < −1, or toward 1 − |ρ| in the case of −1 ≤ ρ < 0. After careful numerical analyses, we found that Eq.(6) can be well approximated by

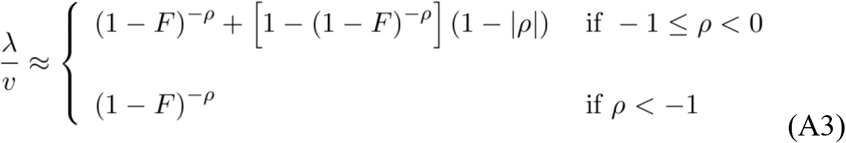

Replacing *ρ* = −*z*|*ρ*_*g*_| in Eq.(A3) and then plugging into Eq.(A2), we have

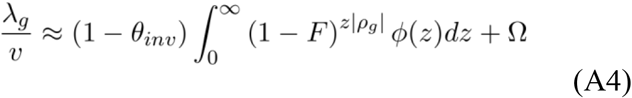

where Ω is given by

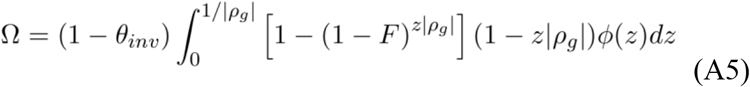

After some calculations, Eq.(A4) turns out to be

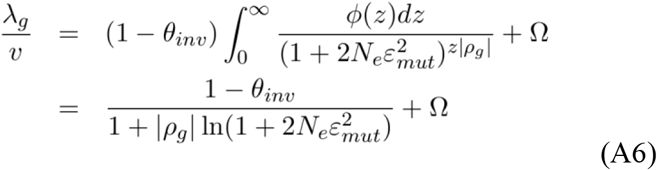

Note that Ω is usually small in practice. When 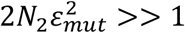, one can show that

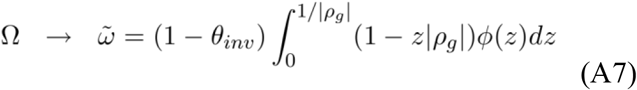

## Acknowledgements

The author is grateful to Andi Clark, Scott Edwards, Nicolas Galtier, Daniel Hartl, Manyuan Long, Jonathan Prichard, and Jeffrey Townsend for their encouragements and constructive comments on the early version of this manuscript.

## Notes

### Competing Interest Statement

The authors have declared no competing interest.

### Summary of Updates

The author has revised the manuscript substantially. The structure of the manuscript follows the Introduction-Methods-Results-Discussion format. A number of typos and grammar/style issues have been corrected. During the last version, I have received many critics and comments, which has been acknowledged appropriately.

## References

Binder BJ, Landman KA, Newgreen DF, Ross JV (2015) Incomplete penetrance: The role of stochasticity in developmental cell colonization. J Theor Biol

Chandler et al. (2013) Does your gene need a background check? How genetic background impacts the analysis of mutations, genes, and evolution. Trends Genet

Chang et al (2008) Transcriptome-wide noise controls lineage choice in mammalian progenitor cells. Nature 453:544–547.

de Jong et al. (2024) Moderating the neutralist–selectionist debate: exactly which propositions are we debating, and which arguments are valid? Biological Reviews

Dowell et al. (2010) Genotype to phenotype: a complex problem. Science 328:469.

Elowitz et al. (2002) Stochastic Gene Expression in a Single Cell. Science

Eldar A et al. (2009) Partial penetrance facilitates developmental evolution in bacteria. Nature 460:510–514.

Ellegren H (2009) A selection model of molecular evolution incorporating the effective population size. Evolution 63:301–305.

Figuet et al. (2016). Life history traits, protein evolution, and the nearly neutral theory in amniotes. Molecular Biology and Evolution

Galtier, N. (2016). Adaptive protein evolution in animals and the effective population size hypothesis. PLoS Genetics.

Galtier (2024) Half a Century of Controversy: The Neutralist/Selectionist Debate in Molecular Evolution. GBE

Gillespie JH (2000) The neutral theory in an infinite population. Gene 261:11–18. 32.

Gillespie JH (2001) Is the population size of a species relevant to its evolution? Evolution 55:2161–2169.

Gu, X (2021) Random penetrance of mutations among individuals: a new type of genetic drift in molecular evolution. Phenomics

Gu, X (2024) Models of fluctuating selection between generations: a solution for the theoretical inconsistency. Journal of Molecular Evolution.

Gu, X (2025) Population genetics and molecular evolution of fluctuating selection among individuals (FSI). BioRxiv.

Hahn MW (2008) Toward a selection theory of molecular evolution. Evolution 62:255–265.

He et al (Bin Su) (2023) Polygenic adaptation leads to a higher reproductive fitness of native Tibetan at high altitude. Current Biology

Hughes, A. L. (2008). Near-neutrality: the leading edge of the neutral theory of molecular evolution. Annals of the New York Academy of Sciences

Hume DA (2000) Probability in transcriptional regulation and its implications for leukocyte differentiation and inducible gene expression. Blood 96:2323–2328

Jensen et al. (2019) The importance of the Neutral Theory in 1968 and 50 years on: A response to Kern and Hahn 2018. Evolution 73:111–114.

Jensen et al. (2025) Genetic modifiers and ascertainment drive variable expressivity of complex disorders. Cell

Karlin and Levikson. 1974. Temporal fluctuations in selection intensities: Case of small population size. Theoret. Popul. Biol. 6: 383–342

Karlin, S, and U. Lieberman, 1974 Random temporal variation in selection intensities: case of large population size. Theoret. Popul. Biol. 6: 355–382

Kern AD, Hahn MW (2018) The Neutral Theory in Light of Natural Selection. Mol Biol Evol 35:1366–1371.

Kimura M (1962) On the probability of fixation of mutant genes in a population. Genetics 47:713–719

Kimura M (1968) Evolutionary rate at the molecular level. Nature 217:624–626.

Kimura M (1979) Model of effectively neutral mutations in which selective constraint is incorporated. PNAS

Kimura M (1983) The Neutral Theory and Molecular Evolution. Cambridge University Press, New York

Khoury MJ, Flanders WD, Beaty TH (1988) Penetrance in the presence of genetic susceptibility to environmental factors.

Kosiol et al. (2008) Patterns of positive selection in six Mammalian genomes. PLoS Genet 4:e1000144.

Lai et al. (2024) Evolution of Phenotypic Variance Provides Insights into the Genetic Basis of Adaptation. GBE

Lanfear R et al (2014) Population size and the rate of evolution. Trends Ecol Evol 29:33–41.

Lee et al. (2022) Evolution and maintenance of phenotypic plasticity. Biosystems.

Lehner B (2013) Genotype to phenotype: lessons from model organisms for human genetics. Nat Rev Genet 14:168–178.

Li WH (1997) Molecular evolution. Sinauer Associates Incorporated, Sunderland, USA Lynch M The Origins of Genome Architecture. Sinauer Associates, Inc, Sunderland, MA

Lynch M et al. (2011) The repatterning of eukaryotic genomes by random genetic drift. Annu Rev Genomics Hum Genet 12:347–366.

Maamar H, Raj A, Dubnau D (2007) Noise in gene expression determines cell fate in Bacillus subtilis. Science 317

McCandlish and Stoltzfus (2014) Modeling Evolution Using the Probability of Fixation: History and Implications. The Quarterly Review of Biology

Mullis et al (2018) The complex underpinnings of genetic background effects. Nat Commun 9:3548.

Munoz-Gomez et al (2021). Constructive neutral evolution 20 years later. JME

Nadeau JH (2001) Modifier genes in mice and humans. Nat Rev Genet 2:165–174.

Nei, M., Suzuki, Y. & Nozawa, M. (2010). The neutral theory of molecular evolution in the genomic era. Annual Review of Genomics and Human Genetics

Ohta, T. (1972). Fixation probability of a mutant influenced by random fluctuation of selection intensity, Genet. Res. Cambridge 19, 33–38.

Ohta T (1973) Slightly deleterious mutant substitutions in evolution. Nature 246:96–98.

Ohta T (1993) An examination of the generation-time effect on molecular evolution. Proc Natl Acad Sci U S A 90:10676–10680.

Ozbudak et al. (2002) Regulation of noise in the expression of a single gene. Nat Genet

Raj A et al. (2010) Variability in gene expression underlies incomplete penetrance. Nature 463:913–918.

Raj A, van Oudenaarden A (2008) Nature, nurture, or chance: stochastic gene expression and its consequences. Cell

Riordan JD, Nadeau JH (2017) From Peas to Disease: Modifier Genes, Network Resilience, and the Genetics of Health. Am J Hum Genet 101:177–191.

Sommer (2020) Phenotypic Plasticity: From Theory and Genetics to Current and Future Challenges. Genetics

Süel (2007) Tunability and noise dependence in differentiation dynamics. Science

Taylor MB, Ehrenreich IM (2014) Genetic interactions involving five or more genes contribute to a complex trait in yeast. PLoS Genet

Vu V et al (2015) Natural Variation in Gene Expression Modulates the Severity of Mutant Phenotypes. Cell

Wagner, A. (2008). Neutralism and selectionism: a network-based reconciliation. Nature Reviews Genetics

Wang et al. (2023) Reproductive variance can drive behavioral dynamics. PNAS

Wernet et al. (2006) Stochastic spineless expression creates the retinal mosaic for colour vision. Nature 440:174–180.

Zuckerkandl E, Pauling L (1965) Evolutionary Divergence and Convergence in Proteins. In: Bryson V, Vogel HJ (eds) Evolving Genes and Proteins. Academic Press, pp 97–166

